# Decoding Angiotensin II Type 1 Receptor Allosteric Communication to Gq and β-arrestin

**DOI:** 10.1101/2021.05.17.444454

**Authors:** Anita K. Nivedha, Yubo Cao, Sangbae Lee, Supriyo Bhattacharya, Stéphane A. Laporte, Nagarajan Vaidehi

**Author notes:** equal contribution (Anita K. Nivedha, Yubo Cao). **Author Contributions** A.K.N. and Y.C. contributed equally to this work. N.V. and S.A.L. conceived the ideas and designed the experiments. A.K.N. and Y.C. executed the work and performed the analysis. S.L. performed MD simulations and some early analysis. S.B. contributed to analysis and discussions. N.V., A.K.N. wrote the paper with edits and suggestions from Y.C. and S.A.L. The authors declare no competing financial interest.

## Abstract

The allosteric communication between the agonist binding site and the G protein or β-arrestin coupling sites in G protein-coupled receptors (GPCRs) play an important role in determining ligand efficacy towards these two signaling pathways and hence the ligand bias. Knowledge of the amino acid residue networks involved in the allosteric communication will aid understanding GPCR signaling and the design of biased ligands. Angiotensin II type I receptor (AT1R) is an ideal model GPCR to study the molecular basis of ligand bias as it has multiple β-arrestin2 and Gq protein biased agonists as well as three-dimensional structures. Using Molecular Dynamics simulations, dynamic allostery analysis, and functional BRET assays, we identified a network of residues involved in allosteric communication from the angiotensin II binding site to the putative Gq coupling sites and another network to the β-arrestin2 coupling sites, with 6 residues common to both pathways located in TM3, TM5 and TM6. Our findings unveil unique and common allosteric communication residue hubs for Gq and β-arr2 coupling by AngII ligands and suggests that some of these residues can be targeted to design biased AT1R ligands. Finally, we show through analysis of the inter-residue distance distributions of the activation microswitches involved in class A GPCR activation for ten different agonists, that these microswitches behave like rheostats with different relative strengths of activation, which we speculate could modulate the relative efficacy of these agonists toward the two signaling pathways.

**Significance Statement:** Knowledge of the residues involved in allosteric communication from the ligand binding site to the G protein or β-arrestin (β-arr) coupling sites in GPCRs will aid in understanding their role in mediating ligand bias. Using a combination of molecular dynamics simulations and functional signaling assays we have identified a network of residues involved in allosteric communication from the Angiotensin II (Ang II) binding site to the Gq and β-arr2 coupling sites in the Ang II type I receptor (AT1R). The residues in the allosteric network for β-arr2 coupling are distributed across multiple structural regions of AT1R compared to Gq coupling. The residues in the two networks show conserved chemical properties across class A GPCRs, demonstrating the importance of allosteric communication in modulating ligand bias.

## Introduction

Angiotensin II (AngII) type 1 receptor (AT1R) belongs to the class A G protein-coupled receptor (GPCR) superfamily of membrane proteins that is activated by the octapeptide hormone Angiotensin II (AngII). AngII is part of the renin-angiotensin-aldosterone system responsible for controlling blood pressure and water retention via smooth muscle contraction and ion transport, and therefore forms an important drug target (1). Binding of AngII to AT1R leads to activation of heterotrimeric Gαq (Gq) family of G proteins, with subsequent engagement of β-arrestin2 (β-arr2). The β-arr2 mediated signaling pathway may lead to pharmacologically desirable cardio protection, while over-activating the Gq coupled pathways leads to undesired cardiovascular effects (2).Therefore, agonists targeting AT1R that preferentially activate the β-arr2 signaling pathway over the Gq pathway commonly known as “biased agonists” are sought after for better therapeutic outcomes (3–9). Understanding the molecular mechanism of biased signaling and the amino acid residues involved in the information flow (or allosteric communication) from the agonist binding site to the Gq and β-arr2 coupling sites would aid the design of biased agonists.

Great strides have been made by recent structural studies on the structure and dynamics of AT1R with different peptide agonists (10–14). The structures of AT1R bound to a nanobody that stabilizes the active state of the receptor and with different biased agonists showed that the receptor adopts very similar conformations in the intracellular regions. However, there were differences observed in certain residue positions in the agonist binding site and in the sodium binding site that is known to be involved in activation of other class A GPCRs (14). An elegant study using double electron-electron resonance (DEER) spectroscopy demonstrated that biased agonists and AngII stabilize conformation ensembles that show distinct differences in the inter-residue distances between labeled residues in the intracellular region of AT1R (10). Single molecule force spectroscopy studies have shown differences in the off rates of the unbinding process of various biased agonists as well as in the number of conformational substates of AT1R when bound to Gq biased agonists compared to β-arr2 biased agonists (11). An all-atom Molecular Dynamics (MD) simulation study by Suomivuori *et al*. on AT1R bound to different biased agonists have shown that AT1R adopts two distinct conformations one of which is speculated to favor β-arr2 coupling and other Gq coupling (13). Using the residue rearrangements in the receptor, they designed and tested a biased agonist. Although these studies provide evidence for ligand specific effects in the intracellular Gq or β-arr2 coupling regions of AT1R, the mechanism of the allosteric communication from the agonist binding region to the Gq and β-arr2 coupling regions remains unclear. Identifying the residues involved in the allosteric communication would provide mechanistic insights into the ligand specific effects on Gq or β-arr2 coupling and thereby the ligand bias.

Here, we used MD simulations for AngII bound AT1R in explicit lipid bilayer, combined with cellular functional BRET assays for measuring AT1R coupling to Gq and β-arr2, to quantitatively predict and test the residue networks that contribute to allosteric communication to Gq and β-arr2 coupling thus giving rise to signaling bias. Our results show that the allosteric communication in AT1R is mediated by a broad network of residues connecting the agonist binding site to the Gq or β-arr2 coupling sites. The network of residues involved in allosteric communication to β-arr2 is spread out across different structural regions (transmembrane helices TM5, TM6, TM7, helix8 and intracellular loop 1 or ICL1) of AT1R compared to the network communicating to the Gq coupling site that involves a majority of residues on TM5 and TM6. We identified six residues that are common to Gq and β-arr2 response (I245^6.40^, S252^6.40^, F249^6.44^, F208^5.5^ H256^6.51^ and R126^3.50^). Despite being common, these residues had differential outcomes on each response when mutated to Ala. F208A^5.51^ showed stronger coupling efficiency to Gq as compared to β-arr2, and as compared to wild type (WT). Although reduced coupling to both pathways compared to WT, F249A^6.44^ also showed stronger coupling efficiency to Gq than β-arr2. On the other hand, R126A^3.50^ exhibited stronger coupling to β-arr2 than Gq compared to WT, while H256A^6.51^ also revealed better coupling efficiency to β-arr2 than to Gq, yet less efficiently in both reponses as compared to WT AT1R.

## Results

Starting from the crystal structure of the partial agonist peptide S1I8 bound active state of AT1R (PDB: 6DO1) (12) we modeled ten different AngII-based peptide agonists, including AngII, SII, six β-arr2 biased, and two G protein biased agonists (shown in Table S1) into this structure. We used the crystal structure of S1I8-AT1R complex structure, because it was the only crystal structure of the active state of AT1R available at the beginning of this study. Follwing 50ns of equilibration procedure for each system (described in the Methods section) we performed five runs each 400ns long of all-atom MD simulations in explicit lipid bilayer (details in Table S2). The resulting aggregated 2μs of trajectories for each of the ten systems were used for further analysis.

### Residues involved in the allosteric communication from AngII binding site to the Gq protein or β-arr2 coupling sites

We previously developed a computational method Allosteer to identify the residues involved in allosteric communication between stipulated sites in a GPCR (15). Allosteer involves calculation of the correlation (or mutual information) in torsion angle distribution for every pair of residues in a given GPCR. For each pair of residues with high mutual information we use graph theory method to calculate the shortest pathway with maximum mutual information. At the end of these calculations, every residue in the GPCR is assigned a “hub score” that quantitfies the number of allosteric communication pathways that pass through that residue (15–18). One could calculate the hub score for all the residues in the user-specified allosteric communication pathways from the agonist binding site to the G protein or β-arrestin coupling sites. We showed that residues located in the allosteric communication pathways modulate the receptor activity depending on the ligand bound to the receptor (15, 17). Here, using *Allosteer* we predicted a set of 26 residues that are shown in Fig. 1A with high hub scores involved in allosteric communication between residues in the AngII binding site and those in the putative Gq protein coupling site. Similarly we predicted a set of 31 residues (Fig. 1C) with high hub scores to β-arr2 coupling site (see Methods for details).

**Figure 1.**
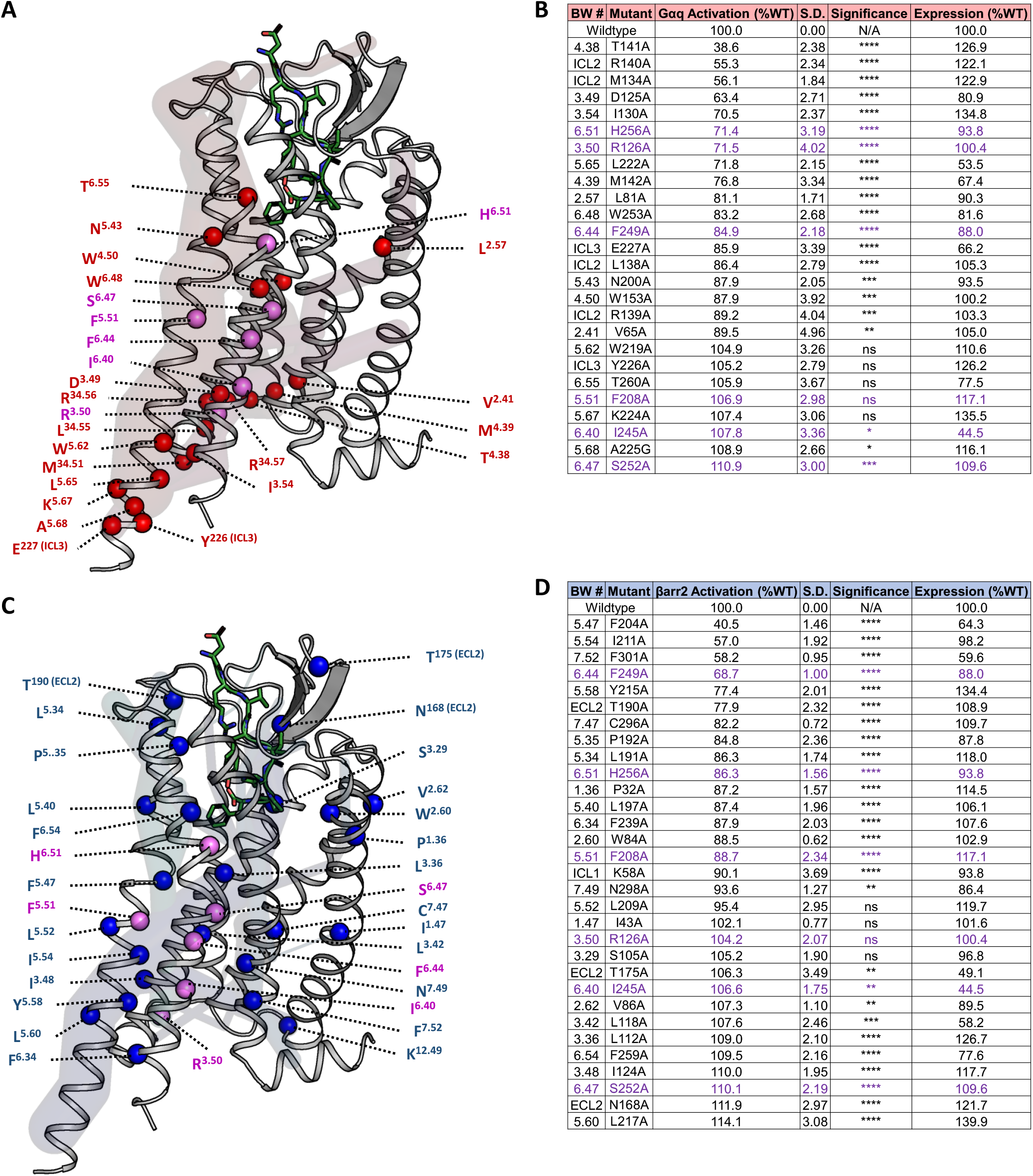
**(A)** AngII (green stick representation) bound AT1R with residues predicted to be involved in allosteric communication to the Gq protein coupling interface (red and purple spheres) that also showed a significant change experimentally in Gq protein coupling upon mutation to alanine using BRET assays. Hub residues are indicated using BW numbering. Residues in purple are hubs common to both Gq protein and βarr2 signaling. The major allosteric communication pipelines housing the hubs contributing to Gq protein signaling are shown in shades of transparent red pipe representation. **(B)** List of allosteric hubs for Gq signaling and BRET measurements on wild type and AT1R mutants expressed with the PKC-c1b sensor in HEK293SL cells and stimulated with various concentrations of AngII to generate concentration-response curves to calculate maximal Gq coupling and effector activation (Emax). Data represent mean ± s.d. from 3 independent experiments, and one-way ANOVA followed by Dunnett’s multiple comparisons test were performed where: **** = p<0.0001, *** = p<0.001, ** = p<0.01, * = p<0.05, n=3. Expression level of receptors determined through whole-cell ELISA is also reported. **(C)** AngII (green stick representation) bound AT1R with residues predicted to be involved in allosteric communication to the β-arr2 coupling interface (blue and purple spheres) that also showed a significant change in β-arr2 coupling upon mutation using BRET assays. The major allosteric communication pipelines to βarr2 interface are shown in shades of transparent blue pipe representation. **(D)** List of allosteric hubs for βarr2 activation and BRET measurements on wild type and AT1R mutants expressed with β-arr2-RlucII and rGFP-CAAX in HEK293SL cells and stimulated with various concentrations of AngII generate concentration-response cuvres to calculate maximal β-arr2 recruitment to receptors (Emax). Data represent mean ± s.d. from 3 independent experiments, and one-way ANOVA followed by Dunnett’s multiple comparisons test were performed where: **** = p<0.0001, *** = p<0.001, ** = p<0.01, * = p<0.05, n=3. Expression level of receptors determined through whole-cell ELISA is also reported.

The residues predicted to be involved in allosteric communication to the Gq or β-arr2 coupled sites were tested for their effect upon mutation to alanine (existing alanine residues were instead replaced with glycines) using BRET-based biosensors in cells (19–22). The AT1R activation of Gq pathway was measured using the downstream PKC BRET biosensor, while β-arr2 engagement was measured using an enhanced bystander BRET sensor, as previously described (19–23). The enhanced bystander BRET assay measures β-arr2 recruitment at the plasma membrane upon receptor activation, and will likely be sensitive to mutations regulating the receptor directly and/or conformationally driven interactions with β-arr2, including those affecting the phosphorylation of receptors (26). We first assessed how receptor expression affects the maximal response of each BRET sensor response by titrating DNA transfection of the wild-type AT1R in cells (see Supplementary text and Figs. S1A to S1D). Next, we expressed wild-type and mutant receptors with BRET biosensors to evaluate their coupling efficacies (E_max_) (Fig. 1B and 1D), as well as the potency of AngII to promote responses in each pathway (EC_50_) (Table S3). All mutant receptors expressed over 50% of WT (except for mutants I245A^6.40^ and T175A^ECL2^) (Figs. 1B and 1D), implying that any loss of signaling responses could not be attributed to a defect in the mutant expression. We evaluated the effect of mutations on receptor activity by comparing their coupling efficacies in each pathway at saturating concentration of AngII where maximal receptor occupancy is reached. This minimizes potential confounding effects of mutations on AngII affinity, which would be mirrored in changes in EC_50_ as seen for some AT1R mutants (Table S3).

Residues that showed a significant change in receptor activity (either positive or negative) for Gq coupling, when mutated, are shown in Figs. 1A and 1B, while those which showed significant changes in β-arr2 recruitment are shown in Figs. 1C and 1D. We next assess quantitatively the performance of our prediction of allosteric hubs using the Receiver Operating Characteristic (ROC) Curve analysis shown in Figs. S3A and S3B. The area under the ROC curve is 0.7 for recovering the residues that affect receptor activity for Gq coupling and 0.6 for βarr2 recruitment, which is a significant enrichment in comparison to random prediction and considering that predictions were made using sparse structural information, independent of experimental data. It is worth noting that that the *Allosteer* method is a system agnostic quantitative method and can be applied to identify allosteric communication networks in any protein without prior knowledge as we have illustrated in these publications(18, 24, 25).

### Allosteric Communication network to the Gq coupling site mainly passes through TM5 and TM6 while to β-arrestin2 spreads across different structural regions of AT1R

The majority of residues in AT1R involved in allosteric communication to the Gq coupling site are located in TM5 and TM6, followed by ICL2, TM3 and TM4 (Fig. 1A, Fig. S4), while residues that communicate to the β-arr2 coupling site are not only more in number, but also distributed more widely across TM5, TM6, TM7, h8 and ICL1 (Fig. 1C, Fig. S4). Analysis of the number of GPCR-G protein contacts in the three dimensional structure of G_q/11_ coupled muscarinic acetyl choline M_1_ receptor (26) shows a lower number of contacts than the GPCR-β-arrestin contacts in the structure of β1-adrenergic receptor-β-arr1 complex (27) (Fig. S5A to S5C). The contacts with β-arr1 are distributed amongst TM5, TM6, ICL2, TM3, TM2, ICL1, TM7 and h8 region of the β1-adrenergic receptor, while those with the G_q/11_ protein are found predominantly in ICL2, TM5, TM6 TM3 and h8 of M_1_ muscarinic receptor. This could be a reason why the residues involved in the allosteric communication to β-arr2 coupling are also spread out across different structural regions of AT1R compared to the residues involved in allosteric communication to Gq.

Analysis of residues involved in allosteric communication network to Gq and β-arr2 in AT1R revealed that six residues located on TM3, TM5 and TM6 are common to both β-arr2 and Gq allosteric communication (Fig. 1). Because we obtained the necessary pharmacological parameters for the signaling effects of these residues when mutated to Ala for both of the two signaling pathways, this gave us the opportunity to determine to what extent their substitution biased the receptors’ responses. When first looking at their coupling efficacies to Gq and β-arr2 (ΔE_max_ Fig. S3C and S3E), we found that mutants F249A^6.44^ and F208A^5.51^ showed significantly more coupling to Gq as compared to β-arr2 in the mutants. F208A^5.51^ showed stronger coupling efficiency to Gq as compared to β-arr2, and as compared to WT. F249A^6.44^ also showed stronger coupling efficiency to Gq than β-arr2 but not compared to both the signaling pathways in the WT. However, H256A^6.51^ and R126A^3.50^ showed the contrary, and I245A^6.40^ and S252A^6.47^ revealed no significant differences between the two signaling pathways. When calculating signaling bias using differences in the relative activity as done previously (21), and despite potential caveats underscored earlier of using such an approach with mutant receptors, we found that while F249A^6.44^ and F208A^5.51^ became more balanced towards both pathways, while H256A^6.51^ and R126A^3.50^ still showed stronger coupling (*e*.*g*. bias) toward β-arr2 compared to Gq in the mutants (Fig. S3D and S3E). R126A^3.50^ exhibited stronger coupling to β-arr2 than Gq compared to WT, while H256A^6.51^ showed weaker responses in both pathways compared to WT AT1R. I245A^6.40^ and S252A^6.40^ remained balanced.These findings suggest that despite these residues belonging to the common Gq and β-arr2 allosteric communication network, some nonetheless differentially contributed to each response, and can be targeted to generate functional selective AT1R. Obviously, other residues we identified at the unique allosteric Gq or β-arr2 hubs may also be targeted by mutagenesis to generate bias signaling AT1R mutants. However, their effects will need to be empirically determined in each complementary pathway to evaluate potential bias. Also, despite identifying key residues for the allosteric Gq and β-arr2 hubs, we cannot exclude the possibility that amino acids elsewhere in the receptor may also have differential bias signaling effects in AT1R when mutated to Ala or other residues. Indeed, in our prediction using *Allosteer* we calculate the hub score for each residue in the wild type AT1R, without considering their substitutions for other amino acids. Further work will be needed to establish the effects of such mutants and their putative biased nature.

### Residues involved in allosteric communication show conserved chemical characteristics across class A GPCRs

To understand the broader functional relevance of the AT1R allosteric residue network to other class A GPCRs, we examined the conservation of their chemical characteristics (see Methods for definitions) among all class A GPCRs (Fig. S6) using the GPCR-SAS webserver (28). 16 out of 26 residues that showed significant change in AT1R coupling to Gq are in the top conserved group of amino acids amongst all class A GPCRs, with 11 of those strictly conserved. Of the remaining 10 allosteric hub residues to Gq coupling, 5 belong to the second most conserved group of amino acids (Fig. S6A). Similarly, 17 out of 31 residues that showed significant change in AT1R coupling to β-arr2 belong to the most conserved group of amino acids in this position, with 8 being strictly conserved. Of the remaining 14 hubs, 8 belong to the second most conserved group of amino acids at this position (Fig. S6B). The evolutionary conservation of the residues involved in allosteric communication from the agonist binding site to the G protein and β-arr2 coupling sites and across all class A GPCRs further underscores the importance of these amino acids for the function of class A GPCRs.

### Relative allosteric communication pipeline strengths account for differences in ligand bias

Recently (17) we advanced the Allosteer method to compute molecular level ligand bias, termed “molecular bias”. We defined the molecular bias as the ratio of strengths of allosteric communication to the Gq protein interface and the β-arr2 interface for a test ligand compared to that of a reference ligand (see Methods). Our prior studies using this method for various biased and reference agonists to β_2_-adrenoceptor, κ-opioid receptor and serotonin receptors demonstrated good correlation with experimentally calculated ligand bias factors (17). We applied the same method to estimate the ligand bias of several agonists to AT1R, using AngII as the reference agonist. The experimental bias factors measured using AngII as the reference agonist were taken from literature (29, 30). As seen in Fig. 2, the calculated molecular ligand bias correlates qualitatively well with the experimental bias factors for both Gq protein biased and β-arr2 biased agonists. This qualitative correlation provides evidence that an atomistic property such as allosteric communication strength is one of the contributing factors that potentiates the ligand bias observed in agonist-receptor pairings. The details of the method used to calculate the ligand bias based on *Allosteer* and caveats of the method are discussed in the Supporting Information.

**Figure 2.**
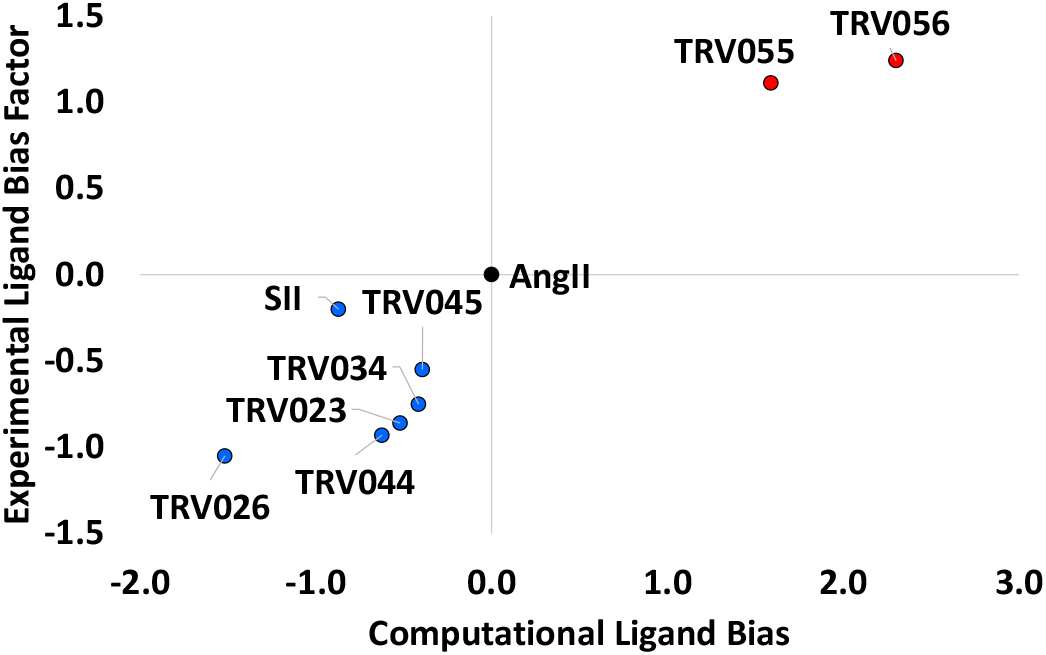
Calculated molecular ligand bias compared to experimental ligand bias factor of Gq or β-arr2 biased agonists for AT1R. The red dots represent Gq biased agonists and the blue dots represent β-arr2 biased agonists. Experimental bias factor values were taken from (29).

### Spatiotemporal heatmap of agonist-AT1R contacts show differences in the binding site features of β-arr2 biased agonists versus Gq biased agonists

Here our goal was to analyze the agonist-specific residue contacts in the binding site of the ten agonist peptides studied here. All the ten agonists are peptide derivatives of the octapeptide AngII and the amino acid positions in the peptides are numbered 0 to 8 (Fig. S7A). The spatiotemporal heatmap in Fig. S7B shows the complete list of agonist-AT1R residue contacts and their persistence calculated from the MD simulation trajectories. The persistence of a ligand-protein residue contact is the percentage of MD snapshots that contain the agonist-receptor contact. The agonist-receptor contacts that show a significant difference in persistence among β-arr2 biased agonists compared to Gq biased agonists or AngII are likely to contribute to the functional selectivity of agonists. For each receptor-agonist contact we calculated the average persistence across the six β-arr2 biased agonists and the same across the two Gq biased agonists. We identified the agonist-AT1R contacts showing a difference in average persistence greater than 10% (which is persistence over a time period of 200ns) as functionally selective to β-arr2 or Gq. As shown in Fig. 3A, the agonist-AT1R contacts that are postulated to contribute to functional selectivity of β-arr2 agonists are mostly located in the N-terminus and TM7, followed by ECL2, TM6 and TM4 with a small percentage of contacts in TM1. On the other hand, the agonist-AT1R contacts that we attribute to functional selectivity of the Gq biased agonists are predominantly located in TM3 and ECL2, followed by TM6, the N-terminus and TM5. 5% of contacts is located in ECL3 and TM7. Residues in TM7 show a difference between β-arr2 biased agonists and Gq biased agonists. Our results could, however, be skewed towards β-arr2 agonists since we have six β-arr2 biased agonists compared to only two Gq biased agonists. Figs. 3B and 3C show the agonist-AT1R residue contacts that show higher persistence in TRV026 or in TRV056 compared to AngII. A large percentage of functional selective contacts for TRV026 compared to AngII are located in TM7 while they come from ECL2, TM5 and TM6 for the Gq biased TRV056. Although these agonist-AT1R contacts are speculated to contribute to functional selectivity, they have not been shown to be causative.

**Figure 3.**
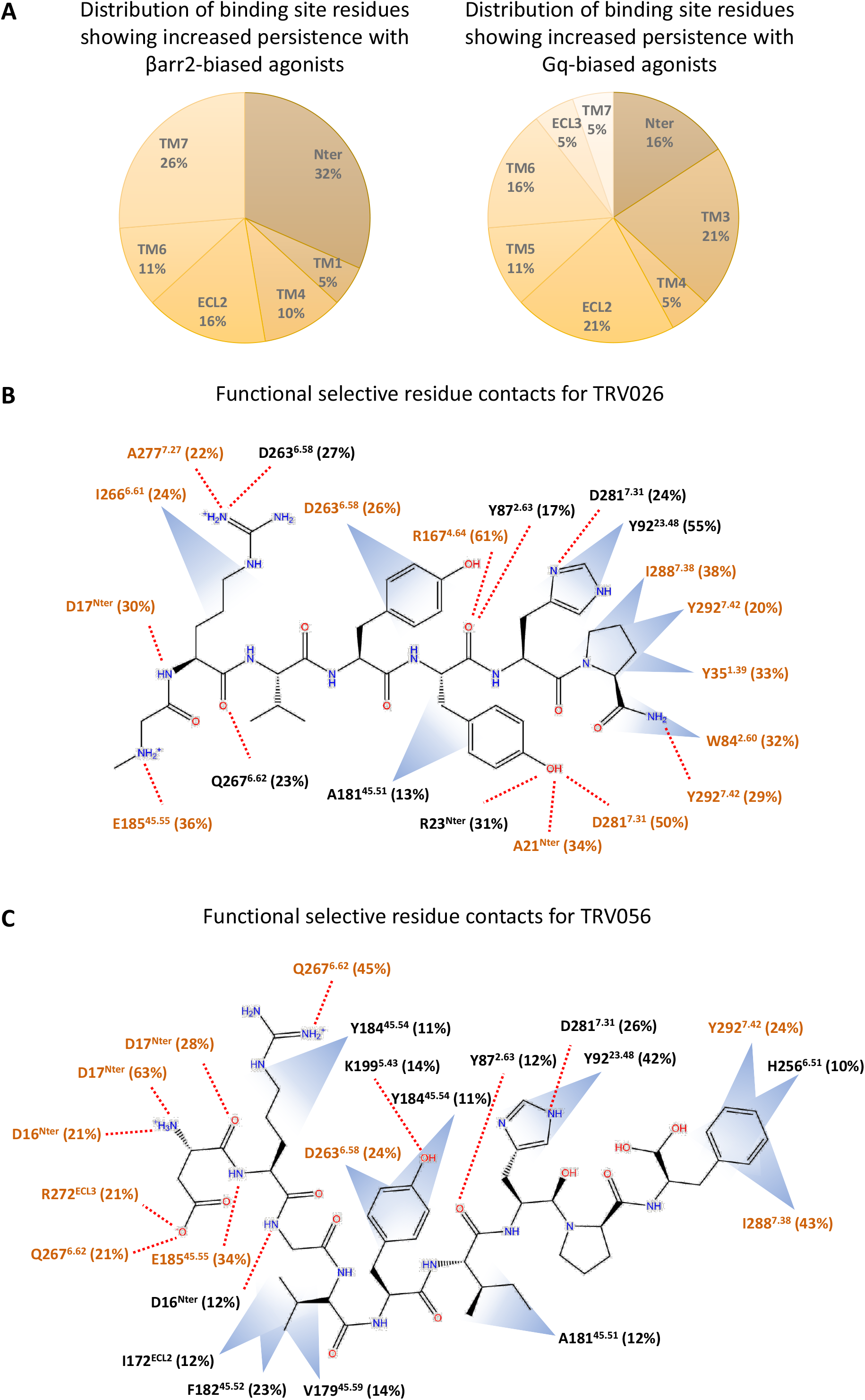
**(A)** Piecharts showing the percentage of agonist-AT1R contacts in each structural region of AT1R. Left: these receptor-agonist contacts show increased (greater than 10% difference which is equivalent to 200ns in persistence) persistence amongst six β-arr2 biased agonists compared to the two Gq-protein biased agonists. Right: these receptor-agonist contacts show increased persistence in the two Gq protein biased agonists compared to the six β-arr2 biased agonists. **(B)** TRV026-AT1R residue contacts that show functional selectivity to TRV026. These residue contacts show increased persistence with TRV026 compared to AngII in AT1R and the difference in persistence is shown in parenthesis. Residue contacts observed exclusively in TRV026-AT1R dynamics and not present in AngII are shown in brown text. Polar contacts are indicated using red dashed lines and Van der Waals contacts are shown in blue shaded triangles. **(C)** TRV056-AT1R residue contacts with increased persistence for Gq biased TRV056 compared to AngII in AT1R. Residue contacts observed only in TRV056-AT1R dynamics and not present in AngII-AT1R dynamics are shown in brown text.

### Activation microswitches behave as rheostats and show differential levels of activation in the presence of different agonists

Comparative analysis of inter-residue distances between inactive and active state structures of class A GPCRs showed significant changes in certain inter-residue distances known as “activation microswitches” (31, 32). Contraction of the inter-residue distances between N/S^3.35^, D^2.50^ and N^7.46^ collectively known as the sodium binding site, distance between P^5.50^ and F^6.44^ in the TM5-TM6 interface and the distance Y^5.58^-Y^7.53^ are three well-characterized “activation microswitches” in class A GPCRs (Fig. 4A). Some of the class A GPCRs show changes in activity towards G protein coupling in response to sodium ion concentration (33). Crystal structures of some class A GPCRs showed the presence of sodium ions in the inactive state structures typically nested between residues S^3.35^, N^3.39^, D^2.50^ and N^7.46^. The sodium ion binding site characterized by these residues shrinks upon receptor activation. However, Wingler *et al*. (14) showed that AT1R has no sensitivity in its activity to sodium ion concentration. Their crystal structure of AngII-AT1R active state complex showed outward movement of N111^3.35^ compared to TRV026 or TRV023 bound AT1R active state structures. This resulted in an expansion of the putative sodium binding site in AngII-AT1R compared to antagonist bound inactive state structure of AT1R or even the TRV026-AT1R active state structure.

**Figure 4.**
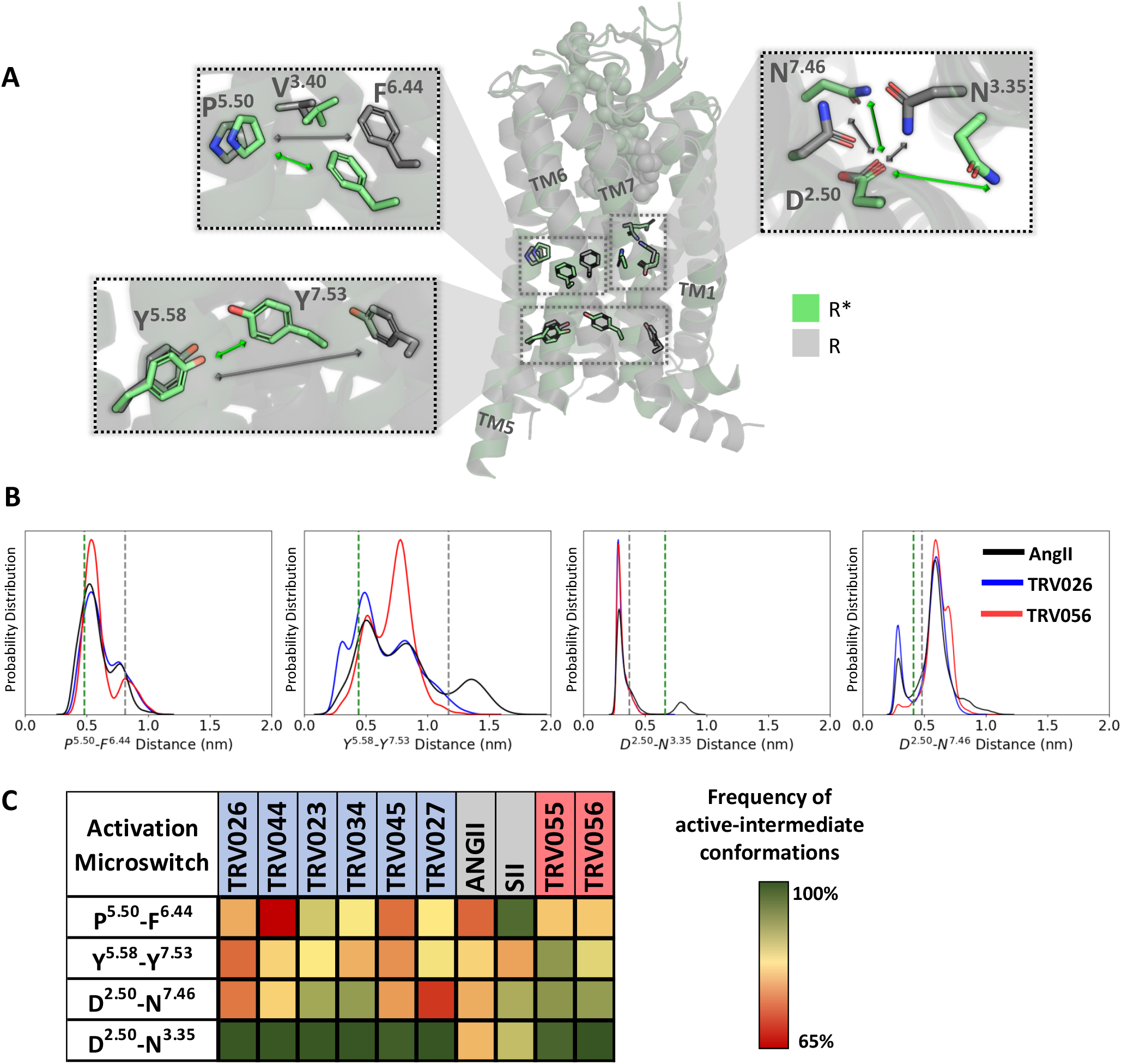
Class A GPCR activation microswitches show distinct patterns of activation with different biased agonists: **(A)** Four inter-residue distances that characterize GPCR activation known as activation microswitches are shown in the central figure. The green cartoon is the nanobody bound active state structure of AngII bound AT1R, R* (PDB: 6OS0) and the grey structure is the antagonist bound inactive state of AT1R, R (PDB: 4YAY). (**B)** The population density distribution of the inter-residue distances in the microswitches during the MD simulations of AT1R with various biased and balanced agonists are shown. The distances measured in AT1R are P207^5.50^ – F249^6.44^, Y215^5.58^-Y302^7.53^, D74^2.50^-N295^7.46^ and D74^2.50^-N111^3.35^. The green and grey dashed lines show the corresponding inter-residue distances in the active (PDB: 6OS0) and inactive (PDB: 4YAY) state crystal structures respectively. **(C)** Heatmap of microswitch distances showing the level of activation for the different agonists when no nanobody or G protein is bound to the receptor, obtained by comparing to reference values obtained from the fully active state AT1R crystal structure (PDB: 6OS0).

Activation microswitches are thought of as “on or off” binary state switches, a concept emerging from the analysis of static structures. Taking the ensemble approach of GPCR conformations, we posited that the activation microswitches for a ligand-receptor pairing could exhibit a rheostat like behavior with different levels of activation in each microswitch (34). To examine the effect of biased ligands on activation microswitches we calculated the inter-residue distance distribution from the 2μs MD simulation trajectories for each of the ten agonists (Figs. 4B and S8). For each microswitch and for each agonist, we also calculated the percentage of MD snapshots (frequency) that move away from distances typified by the endogenous agonist AngII bound active state crystal structure. These frequencies show the percentage of snapshots that are neither in the fully active state or the inactive state (Fig. 4C). All the MD simulations show increased flexibility in the receptor when bound only to an agonist and in the absence of a G protein or nanobody. As a result, in the heatmap shown in Fig. 4C we observed that different agonists show a combination of different levels of occupancy of conformations that are intermediate to active and inactive states of each microswitch despite initiating the MD simulations from the same starting conformation. Our observations imply that the activation microswitches act as rheostats with different levels of activation rather than as binary on or off switches.

AngII-AT1R complex shows the most flexibility in the distance distributions of all microswitches compared to β-arr2 or Gq biased agonists as seen in Fig. 4B and Fig. S8. For example, AngII shows a small peak beyond 11.7Å (distance in the inactive state structure PDB ID: 4YAY) in the Y^5.58^-Y^7.53^ microswitch distance as shown in Fig. 4B. TRV055-AT1R complex also shows similar level of flexibility as AngII in the microswitches (Fig. S8). The β-arr2 biased agonists TRV026, TRV027 and TRV044 show a peak at the contracted D^2.50^-N^7.46^ distance around 3.3Å similar to the distance observed in the active state crystal structures of TRV026-AT1R (3.3Å) and TRV023-AT1R (3.1Å) as opposed to 4.1Å observed in the AngII-AT1R active state structure. There is little variation in the frequency of the D^2.50^-N^3.35^ microswitch across the β-arr2 and Gq protein biased agonists. This could be because for the β-arr2 biased agonists there is little change in the D^2.50^-N^3.35^ distance observed in the active state crystal structure of TRV026-AT1R complex compared to the inactive state crystal structure of AT1R. In the AngII-AT1R complex there is a minor peak in the D^2.50^-N^3.35^ distance distribution that comes from a small conformation ensemble showing outward movement of residue N111^3.35^ in our MD simulations (MD simulations were started from the active state conformation extracted from S1I8:AT1R structure which has the N111^3.35^ facing inside the TM bundle). This leads to the orange cell in the heatmap for AngII (Fig. 4C). In summary, with just the agonist bound, the activation microswitches do not show a definitive pattern of the level of activation of each microswitch among the β-arr2 biased agonists (Fig. 4C). Instead, they show ligand specific combination of the level of activation of microswitches. In addition, the extent of activation of each microswitch is also ligand specific with rheostat-like behavior. This is understandable since the efficacy of these agonists towards β-arr2 coupling arises from an ensemble of conformations with different levels of activation of each of the microswitches.

## Discussion

Allosteric communication from the extracellular agonist binding site to the intracellular core of GPCRs is one of the key molecular factors contributing to ligand efficacy towards G protein and β-arrestin signaling pathways. AT1R is an ideal model GPCR system to study the molecular basis of allosteric communication bias, because it has available, Gq protein and β-arr2 biased agonists with experimentally measured bias factors, and crystal structures of nanobody-bound AT1R active state.

Starting from the nanobody bound AT1R crystal structures, we performed MD simulations in the presence of ten different agonists to extract the resulting structural differences in the receptor. Using a combination of the computational method *Allosteer* on the MD trajectories and BRET functional assays, we have enumerated a network of AT1R residues involved in the allosteric communication from the extracellular AngII binding site to the Gq coupling site and another network to the β-arr2 coupling site. Majority of these residue positions are either conserved or conservative replacements among class A GPCRs showing the functional importance of these positions. While the allosteric residue network to the Gq coupling site are largely localized on TM5 and TM6, those that communicate to the β-arr2 coupling site pass through TM5, TM6, TM7, h8 and ICL1, indicating a wider residue network covering more structural regions of the receptor than for Gq coupling. We speculate that this could be one of the reasons why it is more difficult to tune down β-arr2 mediated signaling than Gq signaling in AT1R (30). Our study showed that mutation of residues F249A^6.44^ and F208A^5.51^ increased coupling efficiency to Gq compared to β-arr2, while H256A^6.51^ and R126A^3.50^ biased coupling to β-arr2 compared to Gq, suggesting that residue positions we identified in the common and specific Gq protein and β-arr2 hubs could be targeted to engineer AT1R biased mutants using alanine and/or non-alanine amino acid substitutions. We defined the molecular level ligand bias as the ratio of strength of allosteric communication to the Gq protein interface and the β-arr2 interface for a test ligand compared to a reference ligand. The molecular bias thus calculated shows a qualitative correlation with experimental bias factors demonstrating that allosteric communication is one of the factors influencing ligand bias. One caveat of our approach is that we translated the G protein and β-arrestin binding site residues from single snapshot structures of other class A GPCRs. Also, our prediction of the residue network using *Allosteer* is based on the MD simulations of the wild type AT1R, without considering their substitutions for other amino acids. This is perhaps why, for instance, we did not capture AT1R mutants such as D74N^2.50^ and N111G^3.35^ which have previously been shown to be biased (35, 36).

The spatiotemporal heatmap of the persistence of the agonist-AT1R contacts for all the ten peptide agonists highlights the presence of AT1R residue contacts that are selectively sampled only by β-arr2 biased agonists and others only by Gq biased agonists. The β-arr2 biased ligand contacts with AT1R that confer functional selectivity are located mainly on TM7. We calculated the distance distributions of the three well characterized activation microswitches, namely the PIF motif on TM5-TM6 interface, sodium binding site and inwards movement of Y302^7.53^. As anticipated AT1R is more flexibile in the MD simulations when bound to an agonist compared to when bound to an antagonist. The three activation microswitches show different levels of activation for the ten agonists as can be anticipated when calculated using an ensemble of conformations, suggesting a rheostat like behavior rather than binary on and off switches.

## Materials and Methods

### Receptor and agonist preparation for AT1R-agonist system setup

All MD simulations of wild type AT1R in the active and inactive states bound to the unbiased, β-arr2 biased and Gq protein biased agonists (Table S1) were performed using the GROMACS package (37) with the CHARMM36 forcefield (38) for proteins, POPC lipids, ions, and using CHARMM TIP3P water as solvent. MD simulations of ten agonists bound to the fully active state of AT1R were performed for AngII, SII, TRV023, TRV026, TRV027, TRV034, TRV044, TRV045, TRV055, and TRV056. The starting structure was taken from the active state structure of S1I8 bound AT1R (PDB: 6DO1) (12). Starting agonist positions for all ten AT1R-agonist complexes were obtained by mutating the corresponding residues back from the native ligand S1I8 in PDB: 6DO1 (see Supplementary Information). The amino acid sequence of all the AngII derivative peptides used in this study and their experimental bias factors are shown in Table S1. The protein structure was prepared and equilibrated for 50ns using procedure described in the Supplementary Information. The equilibration step was followed by 5 production runs with different starting velocities each 400ns long. The snapshots were stored every 20ps and the entire 400ns x 5 runs amounting to 2μs of simulation time per system was used for analysis. Details of the MD simulations and the systems are listed in Table S2. Tests for convergence of the MD simulations were done using overlap between principal components (Fig. S9), methods used to cluster the conformations and other MD analyses are provided in Supplementary information text.

### Method to calculate Allosteric Communication and Ligand Bias

We used the *Allosteer* method (16, 17, 39) to identify the network of residues in the allosteric communication to Gq and β-arr2 coupling sites. Using the aggregated trajectories from MD simulations adding to 1μs for each agonist bound AT1R, we calculated the mutual information in torsion angle distributions for all pairs of residues in the receptor (see Fig. S10 for a flow chart of this procedure). For the residue pairs in the top 10% of the mutual information, we calculated the shortest allosteric communication pathway from the extracellular region of the receptor traversing through the residues in the agonist binding site to the Gq protein or β-arr2 coupling interface residues. See text in Supporting Information for more details. For each residue in AT1R, the number of allosteric communication pathways passing through them is called the “hubscore”. This quantifies the contribution of that residue to the allosteric communication to the β-arr2 and Gq protein interfaces. The residues in the β-arr2 and Gq protein coupling interfaces were taken by translating this information from other solved Gq protein and β-arr2 bound GPCR complex structures. The list of AT1R residues posited to be in the β-arr2 and Gq protein coupling interface used in this study are listed in Table S4. For each residue we calculated the difference in hub score to Gq coupling site and β-arr2 coupling site. We performed quartile analysis of this difference in hub score for identifying Gq protein hubs. As seen in Fig. S2, those residues in the last quartile of this hub score distribution were predicted to have either increased or decreased receptor activity to Gq protein coupling upon mutation. For the prediction of β-arr2 hubs, we used the hub score distribution of all residues to the β-arr2 coupling site and employed the last two quartiles to identify β-arr2 hubs in the AngII-AT1R system. We also used *Allosteer* to calculate the ligand bias as described in the Supplementary Information.

### Inter-residue distance measurements for activation microswitches and generation of heatmap for activation microswitches

The three inter-residue distances D74^2.50^-N111^3.35^, D74^2.50^-N295^7.46^ and N111^3.35^-N295^7.46^ for defining the sodium binding site were calculated as the minimum distance between the side chain heavy atoms of the participating residues, i.e., CB, CG, OD1, OD2 for aspartate, and CB, CG, OD1, ND2 for asparagine. The distance between the Y215^5.58^ and Y302^7.53^ is a microswitch measured between the hydroxyl oxygen atoms, i.e., OH, of the two tyrosine residues. The PF microswitch is measured as the minimum distance between any of the carbon atoms in the side chains of P207^5.50^ i.e., CB, CD, CG and of F249^6.44^ i.e. CB, CG, CD1, CD2, CE1, CE2. These measurements were carried out using the *mindist* module in GROMACS (37). More details are given in the Supporting Information.

### Ligand Receptor contact frequency heatmap generation

The heatmap shown in Fig. S7B were generated for all receptor-ligand contacts using GetContacts (40). The pie charts (Fig. 3A) were constructed by calculating the difference in average persistence frequencies of each contact across all βarr2-biased agonists and all Gq-biased agonists. If the calculated difference in the average frequency of a contact is ≥ 10% for β-arr2 compared to Gq agonists, it has been counted as having a higher persistence frequency with either βarr2-biased agonists or Gq-biased agonists. For Figures 3B and 3C, the contacts in TRV026 or TRV056 that had a higher persistence frequency compared to AngII of ≥10% have been shown, with those residues in brown indicating contacts uniquely formed in TRV026 or TRV056.

### Site-directed receptor mutagenesis

A two-fragment PCR approach was used for the mutagenesis, as described previously (41). Briefly, site-directed mutagenesis primers were generated with 18bp of Gibson homology for Gibson assembly recombination to insert alanine mutations into the signal peptide-FLAG-tagged-AT1R (42). Mutations were made through stepdown PCR, where two separate PCRs were run to split the vector in half. Each half of PCR samples were then purified, and Gibson ligated. The re-annealed vector was transformed into *E. coli* and one colony was picked and amplified. Plasmid DNA was purified using QIAprep Spin Miniprep kit (QIAGEN) and sequenced to verify successful mutagenesis.

### BRET Assay

To measure Gαq or β-arr2 activation, HEK293SL cells were co-transfected with WT or mutant receptors along with either PKC-c1b BRET sensor (19), or β-arr2-RlucII and the plasma membraned-anchored rGFP-CAAX (22), respectively. Cells were transfected and seeded on poly-ornithine-coated, 96-well white plates (BrandTech). 48 hours later, cells were incubated with Tyrodes buffer (140 mM NaCl, 2.7 mM KCl, 1 mM CaCl_2_, 12 mM NaHCO_3_, 5.6 mM D-glucose, 0.5 mM MgCl_2_, 0.37 mM NaH_2_PO_4_, 25 mM HEPES, pH 7.4) at room temperature for 1 hour. Cells were subsequently stimulated with various concentrations of AngII (10^−12^M – 12^-5^M) for 2 min to induce maximal receptor activation and generate concentration-response curves to calculate EC50 and Emax for Gαq or β-arr2 activation. The Rluc substrate coelenterazine 400a (2.5 μM; NanoLight Technology) was added 3 min before BRET measurement. BRET measurements were performed using the Synergy 2 microplate reader (BioTek) with donor filter (410 ± 80 nm) and acceptor filter (515 ± 30 nm). BRET ratios were calculated by dividing the intensity of signal emitted by acceptor over the signal emitted by donor, and BRET change was calculated as the difference between AngII-promoted BRET ratio and unstimulated BRET ratio for each receptor.

### Whole-cell ELISA

To measure cell surface expression of receptors, HEK293SL cells were transfected with WT or mutant receptors and seeded on poly-L-lysine-coated, 96-well clear plates (Fisher Scientific). 48 hours later, cells were washed with PBS and fixed with 4% PFA. Cells were then blocked with 2% BSA and incubated with anti-FLAG M2-peroxidase (Sigma). Cells were washed and colorimetric HRP substrate (SIGMAFAST OPD) was added into each well. After 10 min incubation, 3 M HCL was added to stop the reaction. The plate was then read at an absorbance of 492 nm using Synergy 2 microplate reader (BioTek). To obtain specific signal, non-specific signal from pcDNA-transfected cells was subtracted.

## Acknowledgements

This work was funded by grants from the National Institute of Health (2R01-GM097261) to N.V. and the Canadian Institutes of Health Research (CIHR, MOP-74603) to S.A.L. Y.C. is supported by a doctoral training scholarship from the Fonds de Recherche Santé Quebéc.

